# FetalBoneData: an R data package collating raw measurements of fetal bones across different gestational stages

**DOI:** 10.1101/2025.08.28.672847

**Authors:** Thomas George O’Mahoney, Jael Vakil Kumar

## Abstract

**Objectives:** Raw data of fetal measurements is often difficult to track down in the literature, and researchers are often limited to comparing their own original data to summary tables in synthetic volumes, or investing considerable time and resources into collecting this data themselves. Here, we present the R data package ‘FetalBoneData’, which we hope will improve access to such datasets.

**Methods:** Data was sourced from the literature (primarily Fazekas and Kósa’s (1978), which is long out of print) and work by the lead author. This was collated into a series of.csv files, before being put together into an R data package.

**Results:** We apply the data in this package to compare the measurements of the humerus in a 19th Century fetal collection (Liverpool fetal collection, this paper) against that reported by Fazekas and Kósa (1978) as a case study of the utility of the package.

**Discussion:** The benefit of publishing such data in an open-source format, easily accessible through a popular statistical package, can significantly improve the availability of this type of data. It is hoped that data will be continuously added to the package, further improving its utilization.

## Introduction

Collections of identified fetal remains are rare and hard for many researchers to access. Much data on them, especially raw measurements, are dispersed across the literature in older out of print volumes, making cross comparisons hard for researchers and promoting a reliance on synthetic works.

Much of the literature reports summary statistics only based on data from in-vivo or post-mortem clinical imaging (e.g. (Adalian et al., 2001; Carneiro et al., 2016; Kasraeian et al., 2017; Loughna et al., 2009; Merz et al., 1989; Odita et al., 1982; Queenan et al., 1980; Ravisankar et al., n.d.; Wiśniewski et al., 2019) or summary statistics from direct bone measurements (e.g. (Bareggi et al., 1996, 1994; Felts, 1954; Gardner and Gray, 1970; Matsushita et al., 1995; Scheuer et al., 1980; Scheuer and Black, 2000). While these data sources are of undoubted utility, many modern methods of analysis are multivariate and/or comparative in nature, and require raw data to assure researchers that the data is accurately analysed.

The rise of open source programming within anthropology and especially the popularity of the programming language R, combined with a push to open science, means that there is an impetus to make raw measurements more widely available. This has multiple benefits, including increasing the speed at which junior researchers can conduct research, and removing barriers to researchers who lack access to funding to visit collections.

It is in this spirit, that we are publishing the R package ‘FetalBoneData’ available on Github at https://github.com/Tomomahoney/FetalBoneData and archived on Zenodo https://doi.org/10.5281/zenodo.15805572

## Materials and Methods

The data package currently consists of raw linear measurements from the following sources: (Fazekas and Kósa, 1978; Grzonkowska et al., 2021, 2023). We have avoided reporting means, standard deviation etc., as this hinders the use of more advanced methods of analysis. All data at present has biological sex assigned to nearly every individual, enhancing its utility. Table 1 gives a detailed description of each dataset in the package.

**Table 1.** Detailed description of the data provided in the R package.

| Data name in R | N | Source | Measurements provided |
| --- | --- | --- | --- |
| basicranial_bones | 149 | Fazekas and Kósa (1978) | Body mass in g, Body length in cm; Age (lunar months); sex; Length and width (in mm) of: Sphenoid, greater wing of sphenoid, lesser wing of sphenoid, petrous part, basal occipital, lateral occipital. |
| extremity_dimensions | 138 | Fazekas and Kósa (1978) | Body mass in g, Body length in cm; Age (lunar months); sex; Length and width (in mm) of: Humerus, femur; Length (in mm) of: radius; ulna- tibia; fibula. |
| facial_bones | 138 | Fazekas and Kósa (1978) | Body mass in g, Body length in cm; Age (lunar months); sex; Length and width (in mm) of: Nasal bone, Zygomatic, maxilla, mandible;<br>Length of: palatine,<br>vomer, inferior nasal<br>concha, longset oblique<br>dimension of maxilla,<br>mandible full length. |
| liverpool_lengths | 44 | This paper | Age; Sex; Length in mm<br>(L+R) of: humerus,<br>radius, ulna, femur,<br>tibia, fibula, clavicle. |
| Ribs | 138 | Fazekas and Kósa<br>(1978) | Body mass in g, Body<br>length in cm Age (lunar<br>months); sex; Length of<br>ribs 1-12 |
| shoulder_pelvic | 138 | Fazekas and Kósa<br>(1978) | Body mass in g, Body<br>length in cm Age (lunar<br>months); sex; length<br>(in mm) of: clavicle,<br>scapula, scapular spine,<br>ilium, sciatic notch,<br>pubic bone, width (in<br>mm) of: scapula, ilium,<br>sciatic notch. |
| small_bones | 133 | Fazekas and Kósa<br>(1978) | Body mass in g, Body<br>length in cm Age (lunar<br>months); sex; |
| squamous_occipital | 75 | Grzonkowska et al.<br>(2021) | Sex; crown-rump length<br>(in mm); gestational<br>age in weeks; right and<br>left vertical diameters,<br>Transverse diameter of<br>interparietal par;<br>transverse diameter of<br>the supra-occipital<br>part; projection surface<br>area; volume (mm<sup>3</sup>) |
| squamous_temporal | 37 | Grzonkowska et al.<br>(2023). | Sex; crown-rump length<br>(in mm); gestational<br>age in weeks; right and<br>left: vertical diameter;<br>saggital diameter;<br>projection surface area<br>(mm<sup>2</sup>); volume (mm<sup>3</sup>) |
| vault_bones | 156 | Fazekas and Kósa<br>(1978) | Body mass in g, Body<br>length in cm; Age (lunar<br>months); sex; Length<br>(in mm) of: atlas, axis,<br>first metacarpal,<br>malleus, incus, stapes;<br>width (in mm) of: body,<br>first metacarpal, incus. |

To load the package in R, use the following commands:

\# install.packages(“devtools”)

devtools::install_github(“tomomahoney/FetalBoneData”)

library(FetalBoneData)

This will load the whole data package, and you can then call individual datasets from it to include in analyses.

In this example, we have called the length of the left humerus from the Liverpool Foetal collection and the humeral length from (Fazekas and Kósa, 1978). These are directly comparable, as Fazekas and Kósa reported the left humerus length only for each specimen.

Artificial Intelligence (AI) useage declaration: Microsoft Copilot was used to refine R scripts in this manuscript.

We plotted the distribution of the measurements, both as a combined box/violin plot (fig. 1), and as a scatter plot of age versus humerus length (fig 2.), using the package ggplot2.

**Figure 1.**
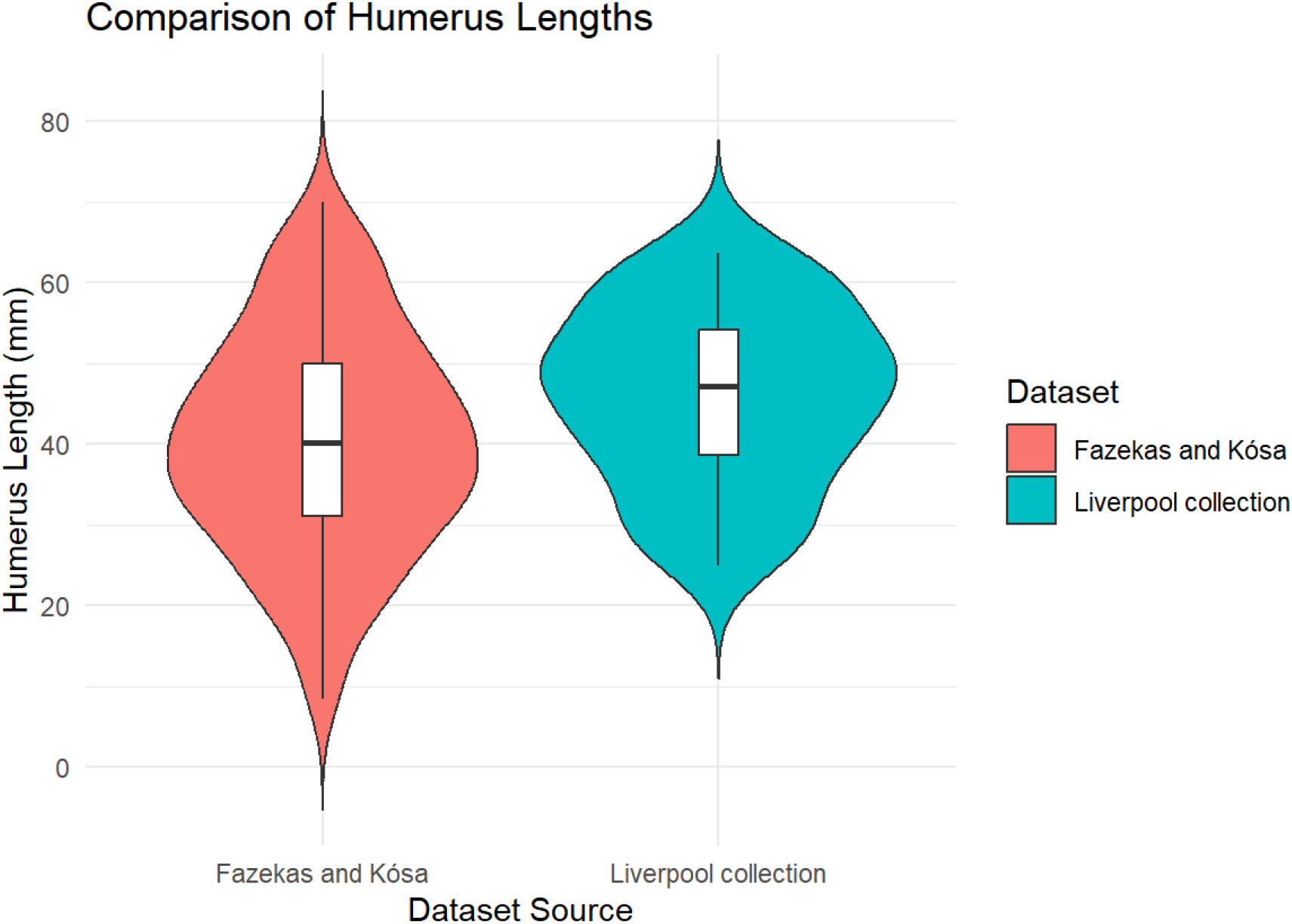
Violin plots of humeral measurement distribution for both datasets.

**Figure 2.**
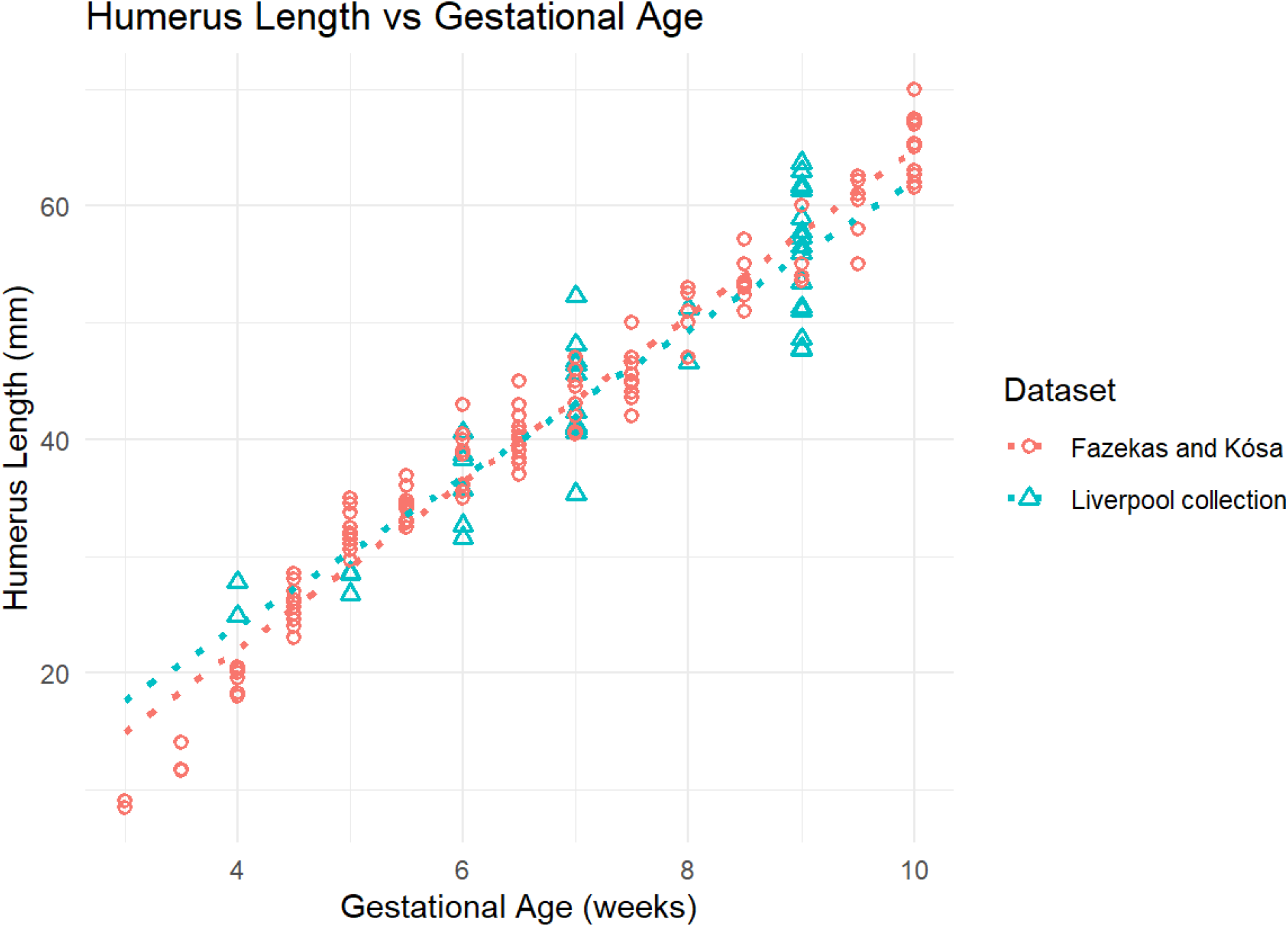
Scatter plot of humerus lengths against gestational ages for each dataset.

We then tested for normality of both datasets using the Shapiro-Wilk test. As data were normally distributed, we subsequently ran Kolmogorov-Smirnov tests to check for distribution differences, as well as a T-test to compare the means. We also displayed the sample distributions using combined violin and box plots and scatter plots.

The code for this is provided in the supplementary information, S1.

## Results

The results were as follows.

## Discussion

In the example above, we can see both from the summary statistics (table 2) and the violin plots (figure 1), combined with the KS test (table 3) that the distribution of measurements is significantly different in the two datasets. This is probably because the measurements from the Liverpool foetal collection only start from 4 months, whereas Fazekas and Kósa reported from 3 lunar months onwards. The t-test (table 3) also confirms a statistically significant difference between the two datasets.

**Table 2.** Summary Statistics of Humerus Lengths.

| Source | Count | Mean | SD | Min | Q1 | Median | Q3 | Max |
| --- | --- | --- | --- | --- | --- | --- | --- | --- |
| Fazekas_Kósa | 138 | 40.24 | 13.74 | 8.5 | 31.1 | 40.15 | 50.00 | 70.0 |
| Liverpool_Collection | 44 | 45.98 | 11.03 | 24.9 | 38.6 | 47.20 | 54.12 | 63.7 |

**Table 3.** Statistical Test Results.

| Test | Statistic | P_value | Normality: Fazekas_Kósa | Normality: Liverpool_Collection |
| --- | --- | --- | --- | --- |
| T-test | -2.821 | 0.006 | 0.408 | 0.132 |
| KS test | 0.250 | 0.022 | 0.408 | 0.132 |

Having access to the raw measurements in this case is undoubtedly an advantage, as we can accurately compare distributions, which can be helpful for assessing trends of secular change (as the two collections are separated by about 100 years). It also demonstrates the speed at which meaningful analyses can be conducted with considerable ease.

To ease updating the package with new data, it is hosted on Github. If any researcher wants to add data to it they can easily do so through GitHub by creating a new branch, adding the data, and submitting a pull request. The pull request is then reviewed and approved if the data are appropriate and submitted correctly. There is a ‘readme’ on the project’s Github page on accepted formats of data. With each version change, the package is also automatically archived on Zenodo, ensuring longevity of availability of the data.

## Conclusion

Here we have presented the new R data package ‘FetalBoneData’ and a simple application of the analysis of the raw data therein. It is hoped that this data package is of utility to biological anthropologists, anatomists and forensic practitioners. It is hoped that over time, this dataset will expand to include more samples, areas of the body, data types, and, gestational ages.

## Supporting information

R scripts for analyses in the article

## Author contributions

**Thomas O’Mahoney:** conceptualization, funding acquisition, data curation, formal analysis, investigation, methodology, software, visualization, writing – original draft, reviewing and editing. **Jael Vakil Kumar:** data curation, investigation, writing-reviewing and editing.

## Acknowledgements

This software package was developed using funding from Anglia Ruskin University Sabbatical fund (awarded to Thomas O’Mahoney).

## Data Availability Statement

All data is freely available on the package’s website:

https://github.com/Tomomahoney/FetalBoneData and archived on Zenodo

https://doi.org/10.5281/zenodo.15805572

All code required to run the analyses is provided as supplementary data to the article.

