## Supplementary material for "FetalBoneData: an R data package collating raw measurements of fetal bones across different gestational stages": R scripts for analyses in the article: FetalBoneData MS supp info.Rmd

````
---
title: "Fetalbonedata MS_supp info"
author: "Tom O'Mahoney"
date: "2025-08-28"
output: html_document
---

```{r setup, include=FALSE}
knitr::opts_chunk$set(echo = TRUE)
```

# Preliminary loading of packages

First, load the necessary packages

```{r}

# Load required libraries
library(ggplot2)
library(dplyr)
library(knitr)
library(FetalBoneData)

```

## Loading of humerus datasets

Loading in the humerus datasets into the R environment

```{r}


# Create data frames
DF1a <- data.frame(
  Gestational_Age = liverpool_lengths$Ag..gstational,
  Humerus_Length = liverpool_lengths$L.Humrus,
  Dataset = "Liverpool collection"
)

DF2a <- data.frame(
  Gestational_Age = Extremity_Dimensions$`lunarmonthold fetuses`,
  Humerus_Length = Extremity_Dimensions$`Humerus Length (mm)`,
  Dataset = "Fazekas and Kósa"
)

# Combine datasets
humerus_age <- rbind(DF1a, DF2a)

```

## Statistical testing

Running summary statistics on the data, and statistical testing for normality and differences between groups

```{r}

# Summary statistics
summary_stats <- humerus_age %>%
  group_by(Dataset) %>%
  summarise(
    Count = sum(!is.na(Humerus_Length)),
    Mean = mean(Humerus_Length, na.rm = TRUE),
    SD = sd(Humerus_Length, na.rm = TRUE),
    Min = min(Humerus_Length, na.rm = TRUE),
    Q1 = quantile(Humerus_Length, 0.25, na.rm = TRUE),
    Median = median(Humerus_Length, na.rm = TRUE),
    Q3 = quantile(Humerus_Length, 0.75, na.rm = TRUE),
    Max = max(Humerus_Length, na.rm = TRUE)
  ) %>%
  mutate(across(where(is.numeric), round, 2))

# Display summary statistics
kable(summary_stats, caption = "Summary Statistics of Humerus Lengths")

# Normality testing
shapiro_df1 <- shapiro.test(DF1a$Humerus_Length)
shapiro_df2 <- shapiro.test(DF2a$Humerus_Length)

normal_df1 <- shapiro_df1$p.value > 0.05
normal_df2 <- shapiro_df2$p.value > 0.05

# Statistical comparison
if (normal_df1 & normal_df2) {
  test_result <- t.test(Humerus_Length ~ Dataset, data = humerus_age)
  test_name <- "T-test"
} else {
  test_result <- wilcox.test(Humerus_Length ~ Dataset, data = humerus_age)
  test_name <- "Wilcoxon test"
}

# Format test results
test_table <- data.frame(
  Test = test_name,
  Statistic = round(test_result$statistic, 3),
  P_value = round(test_result$p.value, 3),
  Normality_Fazekas_Kosa = round(shapiro_df1$p.value, 3),
  Normality_Liverpool_Collection = round(shapiro_df2$p.value, 3)
)

# Display test results
kable(test_table, caption = "Statistical Test Results")

```

## Violin plot

Plotting a combined violin and box plot, with each dataset displayed separately.

```{r}

# Violin plot
ggplot(humerus_age, aes(x = Dataset, y = Humerus_Length, fill = Dataset)) +
  geom_violin(trim = FALSE) +
  geom_boxplot(width = 0.1, fill = "white") +
  labs(title = "Comparison of Humerus Lengths",
       x = "Dataset Source",
       y = "Humerus Length (mm)") +
  theme_minimal()
```

## Scatter plot

Scatter plot with hollow symbols, following the same colourscheme as before.

```{r}
# Plot with hollow symbols and thick outlines
ggplot(humerus_age, aes(x = Gestational_Age, y = Humerus_Length, color = Dataset)) +
  geom_point(aes(shape = Dataset), size = 1.5, stroke = 1, fill = "white") +
  geom_smooth(method = "lm", se = FALSE, linetype = "dotted", size = 1.2, fullrange = TRUE) +
  scale_shape_manual(values = c(1, 2)) +  # 1 = hollow circle, 2 = hollow triangle
  labs(
    title = "Humerus Length vs Gestational Age",
    x = "Gestational Age (weeks)",
    y = "Humerus Length (mm)"
  ) +
  theme_minimal()

```
````
